# Identifying antisense oligonucleotides to disrupt small RNA regulated antibiotic resistance via a cell-free transcription-translation platform

**DOI:** 10.1101/2023.04.19.537543

**Authors:** Min Jen Tsai, Raphael Angelo I. Zambrano, Jeremiah Lyn Susas, Lizette Silva, Melissa K. Takahashi

## Abstract

Bacterial small RNAs (sRNAs) regulate many important physiological processes in cells including antibiotic resistance and virulence genes through base pairing interactions with mRNAs. Antisense oligonucleotides (ASOs) have great potential as therapeutics against bacterial pathogens by targeting sRNAs such as MicF, which regulates outer membrane protein OmpF expression and limits permeability of antibiotics. Here, we devise a cell-free transcription-translation (TX-TL) assay to identify ASO designs that sufficiently sequester MicF. ASOs were then ordered as peptide nucleic acids conjugated to cell-penetrating peptides (CPP-PNA) to allow for effective delivery into bacteria. Subsequent minimum inhibitory concentration (MIC) assays demonstrated that simultaneously targeting the regions of MicF responsible for sequestering the start codon and the Shine-Dalgarno sequence of *ompF* with two different CPP-PNAs synergistically reduced the MIC for a set of antibiotics. This investigation offers a TX-TL based approach to identify novel therapeutic candidates to combat intrinsic sRNA-mediated antibiotic resistance mechanisms.

## INTRODUCTION

The continued increase of antibiotic resistant pathogens is a significant problem in clinical settings and for public health.^1^ Furthermore, bacterial pathogens are acquiring resistance faster than new drugs can be developed.^2,3^ While some resistance mechanisms modify a specific drug or drug target, other mechanisms intrinsic to the bacteria limit the intracellular concentration of antibiotics by reducing membrane permeability and increasing efflux.^4^ These intrinsic mechanisms can broadly affect multiple antibiotics and play a significant role in antibiotic resistance. Specifically, in the presence of antibiotics or other forms of stress, bacteria regulate the production of porins and efflux pumps.^5^ A key component to this regulation are small RNAs (sRNAs) that bind directly to mRNAs and change the accessibility of the Shine-Dalgarno (S-D) sequence or the ability of RNases to degrade the mRNA.^6–8^ The rapid nature of regulation by sRNAs allow bacteria to quickly adapt to the presence of antibiotics.^7,9^ Thus, we propose to develop biomolecules that target sRNAs, preventing their ability to regulate mRNA expression and restore the effectiveness of existing antibiotics.

Antisense oligonucleotides (ASOs) have emerged as an effective method to target both mRNAs and non-coding RNAs in several bacteria species.^10–14^ ASOs are easily designed to target RNAs through complementary Watson-Crick base pairing and several nucleic acid analogues have been developed that are resistant to nuclease activity.^15^ In particular, peptide nucleic acids (PNAs) provide a neutral, peptide-like backbone resistant to both nucleases and proteases.^16^ While PNAs have been shown to penetrate highly permeable bacterial strains,^10^ their delivery is enhanced when conjugated to cell-penetrating peptides (CPPs).^17^ Optimal cellular delivery and RNA target binding has been observed with PNAs 10-12 nucleobases in length.^18,19^ However, there are sRNA-mRNA interactions known to span a longer nucleotide range, on the order of 20-30.^7,20^ Here, we propose the use of synthetic biology methods, namely cell-free transcription-translation (TX-TL) reactions^21^ to identify 10-12 nucleobase long ASOs that are capable of interfering with sRNA regulation of mRNAs. A common method to study sRNA regulation of a specific mRNA target is to create a reporter by fusing the 5’ untranslated region (UTR) and a portion of the gene of interest to the coding sequence of a reporter such as green fluorescent protein (GFP).^22^ GFP expression is then measured in the presence and absence of the sRNA. We extend this method to TX-TL by providing DNA to express the mRNA-reporter, sRNA, and an antisense RNA (asRNA). This enables screening of asRNAs that interfere with sRNA-mRNA regulation.

To demonstrate this approach, we chose the sRNA MicF, which regulates expression of the outer membrane porin OmpF.^20,23^ OmpF is a relatively large porin that bacteria use to take in nutrients especially in low osmolarity conditions.^24^ However, OmpF also allows entry of antibiotics including some β-lactams and quinolones.^5,25,26^ In response to stress, transcription of MicF is activated by one of the transcriptional activators Rob, SoxS, or MarA.^23^ MicF then binds to the 5’ UTR of *ompF* and prevents translation of the porin. In fact, deletion of *micF* from *Escherichia coli* increased susceptibility to some cephalosporins and norfloxacin,^25^ which led to our hypothesis that an ASO designed to sequester MicF could exhibit the same effect.

In this work, we used prior investigations of the MicF-*ompF* interaction to identify potential asRNAs to be screened in TX-TL reactions. We identified a 12-nucleotide long asRNA capable of interfering with MicF-*ompF* binding in TX-TL. However, the CPP-PNA version of the asRNA was unable to increase susceptibility to antibiotics in a minimum inhibitory concentration (MIC) assay. We then proposed a combination PNA approach to target the MicF-*ompF* interaction region more completely. This approach led to a successful increase in susceptibility to cephalothin and nalidixic acid. Furthermore, we validated the use of TX-TL and gene expression assays to identify ASOs that could easily be expanded to other sRNA-mRNA pairs.

## RESULTS AND DISCUSSION

### Initial proof of concept of anti-sRNA strategy

A prior investigation of the interaction between MicF and *ompF* identified that the first 33 nucleotides of MicF bind to *ompF* in the region of the mRNA that encompasses both the S-D sequence and start codon, which suggested that this interaction was responsible for MicF’s regulation of *ompF*.^20^ Therefore, we first sought to test our ability to interfere with the MicF-*ompF* binding by designing an antisense RNA that was complementary to the first 33 nucleotides of MicF. To assess the potential of designed ASOs to interfere with MicF’s ability to repress OmpF expression, we utilized a construct that fused the 5’ UTR and first 13 codons of *ompF* to the coding sequence of superfolder green fluorescent protein (SFGFP).^27^ TX-TL reactions were used to express plasmids containing the fusion reporter and MicF with and without the asRNA anti-MicF 33-1 (Fig 1A). Indeed, anti-MicF 33-1 prevented MicF repression of *ompF* as indicated by the comparable SFGFP fluorescence between the anti-MicF 33-1 test condition and the control expressing the fusion reporter alone (Fig 1B).

**Figure 1.**
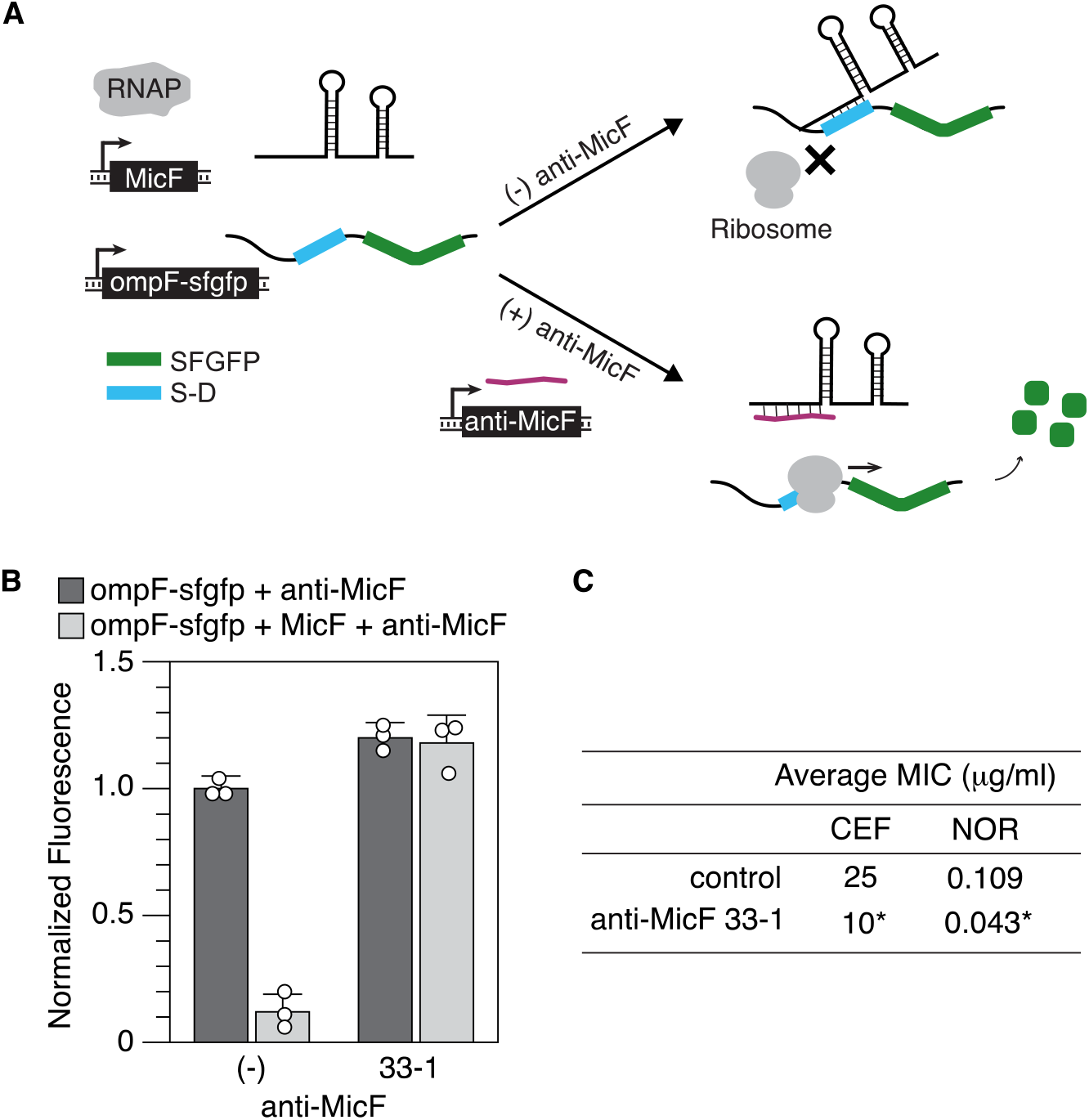
Proof of concept of anti-MicF strategy. (A) Schematic of anti-MicF screening assay. DNA encoding the ompF-sfgfp reporter and MicF are expressed in TX-TL reactions with and without DNA for an anti-MicF asRNA. MicF binds to the 5’ UTR of *ompF* and prevents SFGFP expression. A successful anti-MicF would bind to MicF and prevent it from binding to *ompF*, resulting in high SFGFP expression. (B) Average SFGFP fluorescence from TX-TL reactions testing anti-MicF 33-1. Fluorescence was normalized to reactions containing the ompF-sfgfp reporter alone. Error bars represent standard deviation of three replicates. (C) MIC for cephalothin (CEF) and norfloxacin (NOR) for *E. coli* MG1655 expressing anti-MicF 33-1. Data represents the average of four biological replicates with two technical replicates per biological replicate. Data for each replicate is found in Table S5. * indicates a p-value less than 0.05 for a Wilcoxon Signed-Rank test.

To further support our approach of designing ASOs to target MicF, we then tested if anti-MicF 33-1 could improve antibiotic uptake and thus susceptibility in *E. coli*. To do this, we cloned anti-MicF 33-1 on a plasmid downstream of the inducible promoter p_Lux_.^28^ *E. coli* MG1655 was transformed with the plasmid containing anti-MicF 33-1 or a control plasmid without an asRNA. A broth microdilution MIC assay was performed on both strains for cephalothin and norfloxacin with *N*-acyl homoserine lactone (AHL) added at the time of antibiotic addition to induce anti-MicF production. We observed a decrease in MIC in the presence of anti-MicF 33-1 for both cephalothin and norfloxacin (Fig 1C, Table S5). The reduction in MIC was comparable to that seen in an *E. coli* strain where MicF was deleted from the genome^25^ and thus provided the proof-of-concept for our strategy of targeting MicF with an ASO.

### Identifying the minimal sequence required to inhibit MicF-*ompF* binding

We chose asRNAs to test targeting of MicF because they are easily and affordably cloned into plasmids and expressed in TX-TL reactions. However, to deliver an ASO into cells, it is necessary to use a nucleic acid analog such as PNAs.^15,17^ Furthermore, optimal cellular uptake and RNA targeting occurs with 10-12 nucleobase long PNAs that are conjugated to CPPs.^18,19^ Therefore, we sought to identify a 10-12 nucleotide anti-MicF sequence that could inhibit MicF-*ompF* binding. To do this, we systematically shortened anti-MicF 33-1 from both the 5’ and 3’ ends to identify the regions of MicF important for regulation of OmpF and screened them in TX-TL reactions (Fig 2).

**Figure 2.**
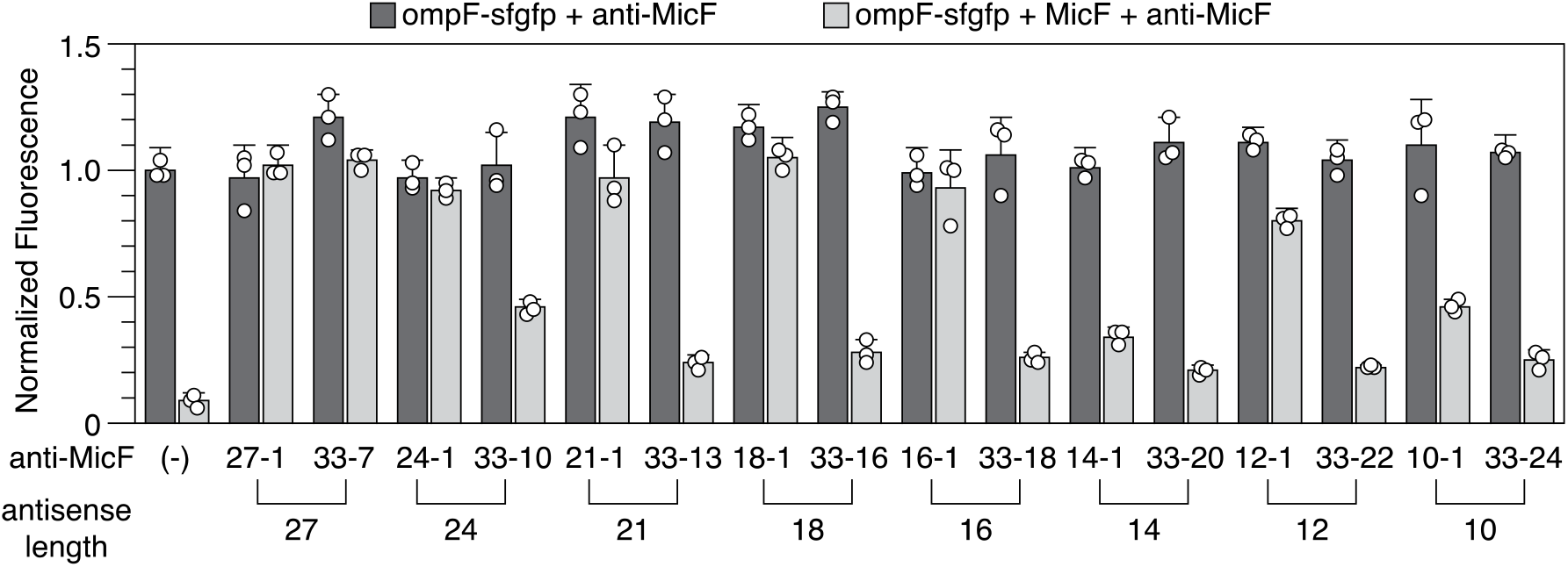
Screening anti-MicF asRNAs. Average normalized SFGFP fluorescence of TX-TL reactions with and without anti-MicFs of various lengths. An anti-MicF is identified by the MicF nucleotides that it was designed to target. Error bars represent standard deviation of three replicates.

Reducing the anti-MicF length from the 5’ end was detrimental and the shortest effective asRNA was 27 nucleotides long (anti-MicF 33-10). In contrast, reducing the anti-MicF length from the 3’ end resulted in a 16-nucleotide asRNA (anti-MicF 16-1) capable of interfering with MicF-*ompF* binding as well as anti-MicF 33-1. Since it did not seem that sequestering the first few nucleotides of MicF (anti-MicF 33-7) was necessary, we sought to reach the goal of a 10-12 nucleotide anti-MicF sequence by creating the anti-MicF constructs 16-5 and 16-7. However, these asRNAs failed to sufficiently interfere with MicF-*ompF* binding (Fig S1). Our best candidate was therefore anti-MicF 12-1, which we then tested in our MIC assay and observed reductions of MIC for both cephalothin and norfloxacin (Table S5).

MicF is predicted to interact with the start codon and S-D sequence of *ompF* at nucleotides 9-11 and 28-31 of MicF, respectively. Our data indicates that interfering with MicF binding to the start codon of *ompF* is key to disrupting regulation. This agrees with prior evidence that the first 13 nucleotides of MicF serve as a seed pairing region and is capable of conferring full regulation of *ompF*.^27^ We tested anti-MicF 13-1 in our TX-TL assay (Fig S1) but did not observe an improvement over anti-MicF 12-1 and thus proceeded with anti-MicF 12-1.

### Assessment of anti-MicF 12-1 in *E. coli*

Next, we ordered anti-MicF 12-1 as a PNA conjugated to the CPP, (KFF)_3_K, which has been used successfully in the past to deliver PNAs into *E. coli*.^14,17,18^ Additionally, we ordered a randomly generated 12-nucleobase scramble PNA to serve as a control (Table S3). We first confirmed that the CPP did not affect the gene expression assay or the ability of the PNA to interfere with MicF-*ompF* binding by testing the CPP-PNA 12-1 and CPP-PNA scramble in TX-TL reactions (Fig S2).

To assess the ability of the CPP-PNA to penetrate cells and bind MicF, we modified the TX-TL gene expression assay to be performed in *E. coli* MG1655. We again used the *ompF-sfgfp* reporter, which was constitutively expressed from a low-copy plasmid and placed MicF under the control of the p_Lux_ promoter on a medium-copy plasmid. After an overnight culture followed by subculture, the CPP-PNA was added together with a low concentration of AHL (2 nM) to induce MicF expression. After one hour of growth, cultures with 6 μM CPP-PNA 12-1 had equivalent SFGFP fluorescence to those with the fusion reporter alone (Fig 3A). Lower concentrations of the CPP-PNA were incapable of interfering with MicF binding as indicated by fluorescence values similar to control cultures with just the fusion reporter and MicF.

**Figure 3.**
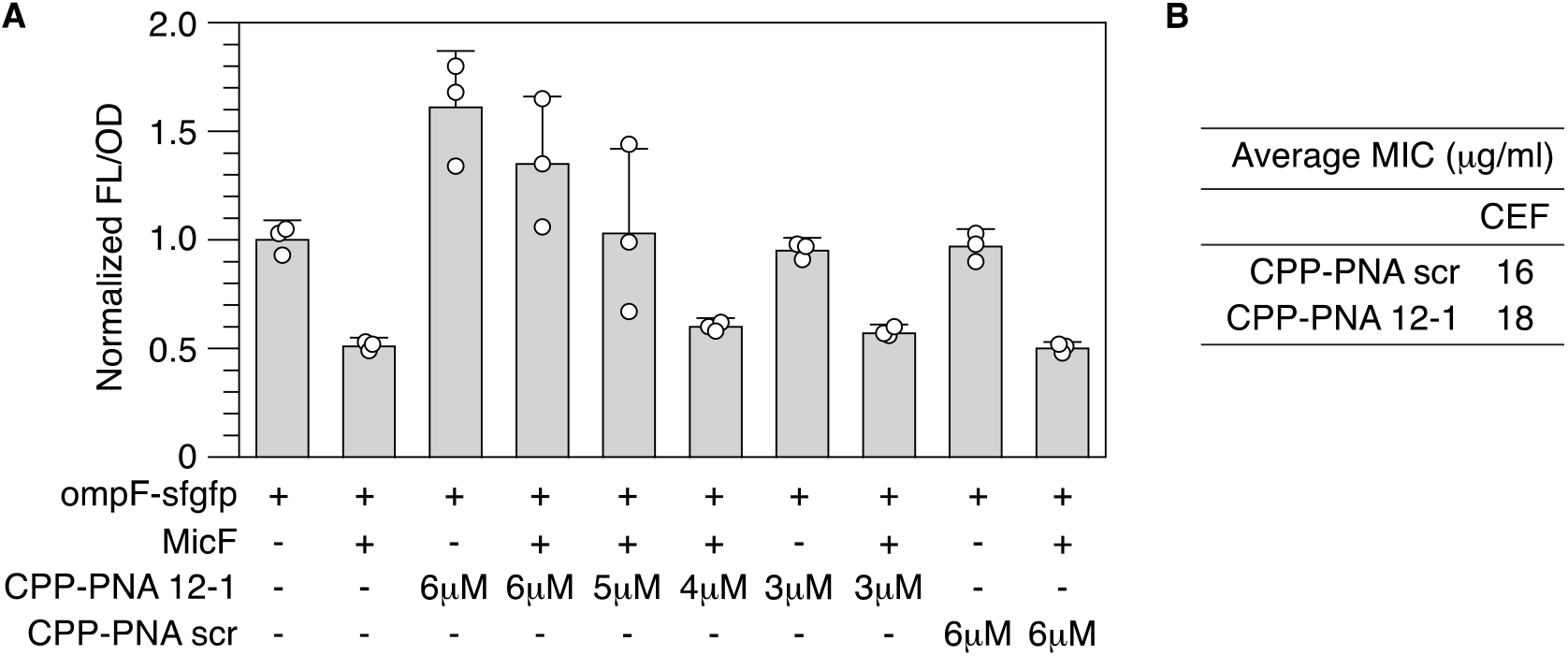
Evaluation of CPP-PNA 12-1. (A) CPP-PNA 12-1 or a CPP-PNA scramble was added to liquid cultures of *E. coli* MG1655 expressing the ompF-sfgfp reporter and an inducible MicF construct. AHL was added at the time of CPP-PNA addition to induce MicF expression. FL/OD was normalized to cells containing the ompF-sfgfp reporter alone. Error bars represent standard deviation of three biological replicates. (B) MIC for cephalothin (CEF) for *E. coli* MG1655 with 6 μM of CPP-PNA 12-1 or 6 μM CPP-PNA scramble. Data represents the average of four biological replicates. Data for each replicate is found in Table S6.

Given the results from our *E. coli* gene expression assay, we then tested the ability of CPP-PNA 12-1 to increase susceptibility to antibiotics. We compared the MIC for cephalothin for *E. coli* MG1655 cultures with either CPP-PNA 12-1 or CPP-PNA scramble added at the time of antibiotics but observed no difference (Fig 3B, Table S6). Although the addition of 6 μM CPP-PNA 12-1 in the *E. coli* gene expression assay led to high SFGFP fluorescence, there was notable variability between the replicates.

Furthermore, when combined only with the fusion reporter, 6 μM CPP-PNA 12-1 exhibited 1.6-fold higher fluorescence than the fusion reporter alone (Fig 3A), which could be confounding the result. The source of the increased fluorescence is unknown, but it could be a concentration dependent effect specific to the PNA 12-1 since 3 μM CPP-PNA 12-1 did not have an effect, nor did 6 μM CPP-PNA scramble.

### A combination of two PNAs to target MicF

To move forward, we proposed utilizing two PNAs that target different regions of MicF simultaneously. We kept PNA 12-1 due to the known importance of the first 13 nucleotides of MicF for its regulation of *ompF*.^27^ We hypothesized that we could use a lower concentration of PNA 12-1 if in combination with a second PNA to avoid the previously-observed concentration dependent effect. For the second PNA, we proposed targeting the nucleotides of MicF that bind the S-D sequence of *ompF* as this was identified as a secondary binding site between MicF and *ompF* through RNA structure probing.^20^ Anti-MicF 33-22 essentially showed no ability to interfere with MicF-*ompF* binding on its own in TX-TL reactions. However, when tested in combination with anti-MicF 12-1 there was an improvement in MicF-*ompF* binding inhibition over anti-MicF 12-1 alone (Fig 4A). Given this finding, we ordered the CPP-PNA 33-22 and confirmed that the combination with CPP-PNA 12-1 effectively allowed *ompF-sfgfp* expression in the TX-TL assay (Fig S3).

**Figure 4.**
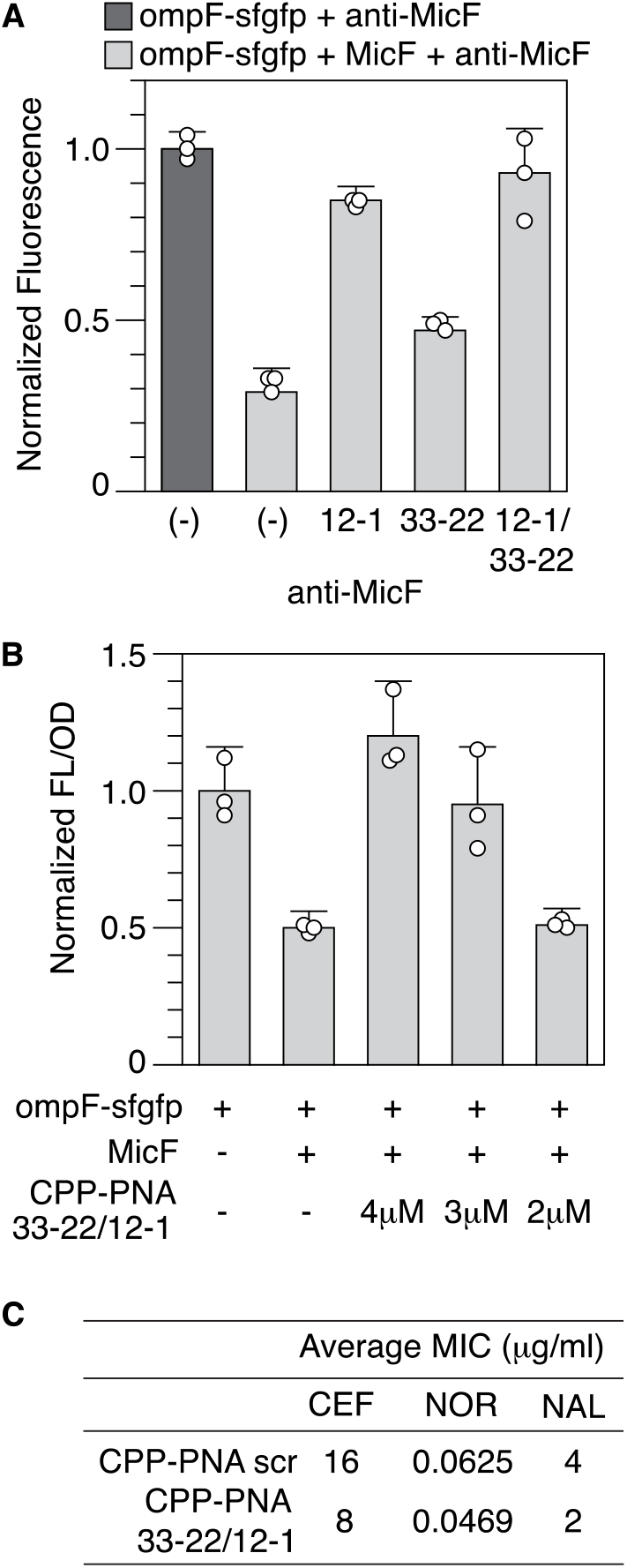
Evaluation of the combination PNA strategy to target MicF. (A) Average normalized SFGFP fluorescence of TX-TL reactions with anti-MicF 12-1 and anti-MicF 33-22 individually and in combination. Error bars represent standard deviation of three replicates. (B) CPP-PNA 12-1 and CPP-PNA 33-22 or a CPP-PNA scramble was added to liquid cultures of *E. coli* MG1655 expressing the ompF-sfgfp reporter and an inducible MicF construct. AHL was added at the time of CPP-PNA addition to induce MicF expression. FL/OD was normalized to cells containing the ompF-sfgfp reporter alone. Error bars represent standard deviation of three biological replicates. (C) MIC for cephalothin (CEF), norfloxacin (NOR), and nalidixic acid (NAL) for *E. coli* MG1655 with 3 μM of CPP-PNA 12-1 plus 3 μM of CPP-PNA 33-22 or 6 μM CPP-PNA scramble. Data represents the average of four biological replicates. Data for each replicate is found in Table S6.

Next, we used our *E. coli* gene expression assay to test different concentrations of the CPP-PNA 12-1 and CPP-PNA 33-22 combination. The data suggested that both 4 μM and 3 μM of each CPP-PNA could inhibit MicF-*ompF* binding in *E. coli* (Fig 4B, Fig S4). However, when examining the control condition that introduced 8 μM of the CPP-PNA scramble, we observed a decrease in growth rate of the cells when compared to 6 μM of the CPP-PNA scramble and *E. coli* MG1655 alone (Fig S5A). To confirm this result, we performed an MIC for cephalothin comparing 8 μM and 6 μM CPP-PNA scramble and indeed observed a reduction in MIC for the 8 μM CPP-PNA scramble, which indicates that the CPP-PNA at 8 μM is inhibiting growth of *E. coli* (Fig S5B). Therefore, to assess the effect of our specific PNA sequences we moved forward with the 3 μM combination of CPP-PNA 12-1 and 33-22.

Finally, we tested if the combination CPP-PNAs could improve susceptibility to antibiotics known to use OmpF for entry into cells. A consistent reduction in MIC was observed for cephalothin, but only half of the biological replicates for norfloxacin resulted in a reduction in MIC (Fig 4C, Table S6). Since the effect on norfloxacin was not significant, we performed an MIC for nalidixic acid, which belongs to the same quinolone family of antibiotics as norfloxacin and observed a consistent reduction in MIC using our combination CPP-PNA 12-1 and 33-22.

## CONCLUSION

In this work, we demonstrated that targeting sRNAs with ASOs to counteract an intrinsic antibiotic resistance mechanism is feasible. Specifically, we designed two PNAs to bind to the sRNA MicF and prevent it from binding to *ompF* mRNA. Addition of the two CPP-PNAs to liquid cultures of *E. coli* led to a decrease in MIC for both cephalothin and nalidixic acid. Theoretically, this is due to the increased influx of antibiotics through continued OmpF porin expression caused by the PNAs and thus will help prolong the usability of existing antibiotics. Our work, and recent work that used ASOs to target the Qrr sRNA of *Vibrio cholerae*,^13^ highlight sRNAs as potential new targets for antimicrobial therapeutics and the use of synthetic biology tools to identify ASOs. The *in vitro* interaction assay developed by Henderson *et al*. allows for identification of ASOs that interfere with sRNA-mRNA binding. An advantage of using an mRNA-sfgfp fusion reporter in TX-TL to screen ASOs is the ability to identify the disruption of the sRNA-mRNA regulation mechanism, which is the goal when targeting sRNAs. Although our *E. coli* gene expression assay was useful for identifying appropriate concentrations of CPP-PNA to use, the TX-TL assay was a better indicator of ASO ability to disrupt MicF-*ompF* regulation. While a single PNA may be suitable to sequester some sRNAs, our findings demonstrate that this was not possible for MicF. Again, the TX-TL assay was capable of screening the combination strategy as we did see improvement using the combination anti-MicF 12-1 and 33-22 over just anti-MicF 12-1 alone. By applying the approach demonstrated here to other sRNAs involved in regulation of antibiotic resistance and virulence, the TX-TL platform could be a powerful tool in the fight against antibiotic resistant pathogens.

## METHODS

### Plasmid construction and purification

All plasmids used in this study are listed in Supporting Information Table S4, with key sequences found in Table S1 and all anti-MicF sequences in Table S2. Plasmids were constructed using inverse PCR or Gibson Assembly. Plasmids for TX-TL experiments were purified using a Qiagen QIAfilter Plasmid Midi Kit (catalog no. 12243) followed by isopropyl alcohol precipitation and elution with nuclease free water. Plasmids for *E. coli* gene expression or MIC analysis were purified using Qiagen QIAprep Spin Miniprep Kit (catalog no. 27104).

### PNA storage and handling

PNAs conjugated to the CPP (KFF)_3_K were ordered from PNA Bio Inc. (Newbury Park, CA). Specific sequences are listed in Table S3. CPP-PNAs were handled per guidance from PNA Bio: lyophilized pellets were thawed to room temperature, protected from light, reconstituted to 250 μM using sterile nuclease free water, and heated at 50°C for 10 minutes. Aliquots were stored at −80°C. Frozen stock solutions were thawed on ice and heated at 50°C for 5 minutes prior to use.

### TX-TL experiments

The myTXTL Sigma 70 Master Mix was purchased from Arbor Biosciences (catalog no. 507005). Reactions were prepared on ice according to manufacturer’s protocol with the addition of 2mM magnesium glutamate (Sigma-Aldrich 49605). Plasmids were provided at the following concentrations: ompF-sfgfp (0.031 nM), MicF (2 nM), and anti-MicF (1 nM). CPP-PNAs were provided at 2 μM. A control plasmid (MKT046) containing the same constitutive promoter and transcription terminator was used to maintain a consistent total DNA concentration in each of the reactions. Three 10 μL aliquots of each reaction mixture were transferred to a 384-well plate (Thermo Scientific 142761), covered with a seal (VWR 60941-078), spun down at 1500 rpm for 30 seconds (Eppendorf 5810R centrifuge), and placed on a Biotek SynergyH1 plate reader. The temperature was controlled at 37°C and SFGFP fluorescence was measured from the bottom of the plate every five minutes (485 nm excitation, 520 nm emission, gain 80).

Data shown is from the 2-hour time point. All fluorescence values were normalized to the ompF-sfgfp alone condition.

### MIC assay – inducible anti-MicF expression

Plasmids containing anti-MicF sequences or a control downstream of the inducible promoter p_Lux_ were transformed into *E. coli* MG1655, plated on LB (BD 244610) + Agar (BD 214530) plates containing 50 μg/mL kanamycin, and incubated overnight at 37°C. Colonies from each condition were inoculated into 5 mL of MOPS EZ Rich Defined Medium (Teknova M2105) containing 50 μg/mL kanamycin and grown for 17 hours overnight at 37°C while shaking at 275 rpm (SHEL LAB SSI3). Cell pellets of the overnight cultures were collected by centrifugation of 250 μL of the culture at 13,000 rpm for 1 minute. The pellets were resuspended in 1 mL of fresh MOPS media (without selective antibiotics) and OD600 was measured using the Biotek Synergy H1 plate reader and 96-well plate (Corning 3370). Cultures were diluted appropriately in MOPS media to reach a final concentration of 5×10^5^ CFU/mL in each well of a 96-well plate (Corning 3370). AHL (Cayman Chemical 143537-62-6) was added to each well at a final concentration of 10 nM. A two-fold serial dilution of the antibiotic was made across the columns of the plate. Cephalothin (Sigma-Aldrich C4520) concentrations ranged from 512 μg/mL to 1 μg/mL. Norfloxacin (Sigma-Aldrich N9890) concentrations ranged from 1 μg/mL to 0.0019 μg/mL. No antibiotic and media only controls were added to additional columns on the plate. Plates were sealed first with a breath-easier membrane (USA Scientific, 9126-2100) followed by a breathe-easy membrane (Diversified Biotech, BEM-1) and incubated for 24 hours at 37°C while shaking at 100 rpm (Labnet Vortemp 56). After incubation, OD600 was measured using the Biotek Synergy H1 plate reader and MIC was determined. Four biological replicates with two technical replicates were performed for each condition.

### *E. coli* gene expression assay

Plasmids containing the ompF-sfgfp reporter, p_Lux_-MicF, or controls were transformed into *E. coli* MG1655, plated onto LB + Agar plates containing 34 μg/mL chloramphenicol and 100 μg/mL carbenicillin, and incubated overnight at 37°C. Three colonies from each condition were inoculated into 300 μL of LB with corresponding antibiotics in a 2 mL 96-well block (Costar 3961) sealed with a breath-easier membrane and grown for 17 hours overnight at 37°C while shaking at 100 rpm (Labnet Vortemp 56). Four microliters of the overnight culture were added to 296 μL of MOPS media that was pre-warmed at 37°C for 30 minutes. The cultures were grown under the same conditions as above for approximately 2.5-3 hours until an OD600 of 0.35-0.4 was reached. Cultures were diluted a second time using pre-warmed MOPS into a 96-well plate (Corning 3631) to reach an OD600 of approximately 0.015. AHL was added to each well at a final concentration of 2 nM. CPP-PNAs were added to each well at final concentrations indicated in the figures. The plate was sealed with a breathe-easy membrane and placed on a Biotek Synergy H1 plate reader. The temperature was controlled at 37°C with continuous double-orbital shaking (425 cpm) for two hours. OD600 and SFGFP fluorescence was measured from the bottom of the plate every ten minutes (485 nm excitation, 520 nm emission, gain 80).

### MIC assay – CPP-PNA

*E. coli* MG1655 was struck out on LB + Agar plates to isolate single colonies. Colonies were inoculated into 5 mL of MOPS media and grown for 17 hours overnight at 37°C at 275 rpm. Cell pellets of the overnight cultures were collected by centrifugation of 250 μL of the culture at 13,000 rpm for 1 minute. The pellets were resuspended in 1 mL of fresh MOPS media and OD600 was measured. Cultures were diluted appropriately in MOPS media to reach a final concentration of 5×10^5^ CFU/mL in each well of a 96-well plate (Corning 3370). A two-fold serial dilution of the antibiotic was made across the columns of the plate. Cephalothin concentrations ranged from 512 μg/mL to 1 μg/mL. Norfloxacin concentrations ranged from 1 μg/mL to 0.0019 μg/mL. Nalidixic acid (Thermo Scientific J63550.06) concentrations ranged from 32 μg/mL to 0.25 μg/mL. CPP-PNAs were added to each well of the plate at final concentrations indicated in the figures. No antibiotic and media only controls were added to additional columns on the plate. Plates were sealed first with a breath-easier membrane followed by a breathe-easy membrane and incubated for 24 hours at 37°C while shaking at 100 rpm. After incubation, OD600 was measured using the Biotek Synergy H1 plate reader and MIC was determined. Four biological replicates were performed for each condition.

## ACKNOWLEDGEMENTS

Research reported in this work was supported by the National Institute of General Medical Sciences of the National Institutes of Health under award number SC2GM136500. R.A.I.Z. was supported by the National Institute of General Medical Sciences of the National Institutes of Health under award number TL4GM118977. The content is solely the responsibility of the authors and does not necessarily represent the official views of the National Institutes of Health.

