## Supplemental Information for "Identifying antisense oligonucleotides to disrupt small RNA regulated antibiotic resistance via a cell-free transcription-translation platform"

Table of contents:


| | **Description** | **Page** |
|---|---|---|
| Table S1 | Important DNA sequences | 2 |
| Table S2 | anti-MicF sequences used in this study | 3 |
| Table S3 | CPP-PNA sequences | 4 |
| Table S4 | Plasmids used in this study | 4-5 |
| Table S5 | pLux-anti-MicF MIC values of individual replicates in μg/ml | 6 |
| Table S6 | CPP-PNA MIC values of individual replicates in μg/ml | 7 |
| Figure S1 | Screening anti-MicF asRNAs (continued) | 8 |
| Figure S2 | TX-TL test of CPP-PNA 12-1 and scramble | 9 |
| Figure S3 | TX-TL test of CPP-PNA 33-22 alone and combination | 9 |
| Figure S4 | Controls for *E. coli* gene expression assay assessing CPP-PNAs | 10 |
| Figure S5 | Effect of CPP-PNA concentration on growth of *E. coli* | 11 |

Table S1: Important DNA sequences

| Name | Sequence |
|---|---|
| pLux promoter | acctgtaggatcgtacaggtttacgcaagaaaatggtttgttatagtcg<br>aataaa |
| LuxR sequence | atgaaaaacataaatgccgacgacacatacagaataattaataaa<br>attaaagcttgtagaagcaataatgatattaatcaatgcttatctgatat<br>gactaaaatggtacattgtgaatattatttactcgcgatcatttatcctca<br>ttctatggttaaatctgatatttcaatcctagataattaccctaaaaaatg<br>gaggcaatattatgatgacgctaatttaataaaatatgatcctatagta<br>gattattctaactccaatcattcaccaattaattggaatatatttgaaaa<br>caatgctgtaaataaaaaatctccaaatgtaattaaagaagcgaaa<br>acatcaggtcttatcactgggtttagtttccctattcatacggctaacaat<br>ggcttcggaatgcttagttttgcacattcagaaaaagacaactatata<br>gatagtttatttttacatgcgtgtatgaacataccattaattgttccttctct<br>agttgataattatcgaaaaataaatatagcaaataataaatcaaaca<br>acgatttaaccaaaagagaaaaagaatgtttagcgtgggcatgcg<br>aaggaaaaagctcttgggatatttcaaaaatattaggttgcagtgag<br>cgtactgtcactttccatttaaccaatgcgcaaatgaaactcaataca<br>acaaaccgctgccaaagtatttctaaagcaattttaacaggagcaat<br>tgattgcccatactttaaaaattaa |
| J23119 promoter | ttgacagctagctcagtcctaggtataatactagt |
| J23118 promoter | ttgacggctagctcagtcctaggtattgtgctagc |
| ompF-sfgfp fusion | agacacataaagacaccaaactctcatcaatagttccgtaaattttta<br>ttgacagaacttattgacggcagtggcaggtgtcataaaaaaaacc<br>atgagggtaataaataatgatgaagcgcaatattctggcagtgatcg<br>tccctgctagcaaaggagaagaacttttcactggagttgtcccaattct<br>tgttgaattagatggtgatgttaatgggcacaaattttctgtccgtggag<br>agggtgaaggtgatgctacaaacggaaaactcacccttaaatttattt<br>gcactactggaaaactacctgttccgtggccaacacttgtcactactc<br>tgacctatggtgttcaatgcttttcccgttatccggatcacatgaaacgg<br>catgactttttcaagagtgccatgcccgaaggttatgtacaggaacg<br>cactatatctttcaaagatgacgggacctacaagacgcgtgctgaa<br>gtcaagtttgaaggtgatacccttgttaatcgtatcgagttaaagggta<br>ttgattttaaagaagatggaaacattcttggacacaaactcgagtaca<br>actttaactcacacaatgtatacatcacggcagacaaacaaaaga<br>atggaatcaaagctaacttcaaaattcgccacaacgttgaagatggt<br>tccgttcaactagcagaccattatcaacaaaatactccaattggcgat<br>ggccctgtccttttaccagacaaccattacctgtcgacacaatctgtcc<br>tttcgaaagatcccaacgaaaagcgtgaccacatggtccttcttgagt<br>ttgtaactgctgctgggattacacatggcatggatgagctctacaaa |
| MicF | gctatcatcattaactttatttattaccgtcattcatttctgaatgtctgttta<br>cccctatttcaaccggatgcctcgcattcggtttttttt |
| t500 terminator | caaagcccgccgaaaggcgggctttttttt |
| T1 terminator | gcatcaaataaaacgaaaggctcagtcgaaagactgggcctttcgt<br>tttatctgttgtttgtcggtgaacgctctcctgagtaggacaaatccgcc<br>gccctaga |

Table S2: anti-MicF sequences used in this study

| Name | Sequence |
|---|---|
| anti-MicF 50-1 | cagacattcagaaatgaatgacggtaataaataaagttaatgatgatagc |
| anti-MicF 33-1 | atgacggtaataaataaagttaatgatgatagc |
| anti-MicF 27-1 | gtaataaataaagttaatgatgatagc |
| anti-MicF 24-1 | ataaataaagttaatgatgatagc |
| anti-MicF 21-1 | aataaagttaatgatgatagc |
| anti-MicF 18-1 | aaagttaatgatgatagc |
| anti-MicF 16-1 | agttaatgatgatagc |
| anti-MicF 14-1 | ttaatgatgatagc |
| anti-MicF 13-1 | taatgatgatagc |
| anti-MicF 12-1 | aatgatgatagc |
| anti-MicF 11-1 | atgatgatagc |
| anti-MicF 10-1 | tgatgatagc |
| anti-MicF 33-7 | atgacggtaataaataaagttaatgat |
| anti-MicF 33-10 | atgacggtaataaataaagttaat |
| anti-MicF 33-13 | atgacggtaataaataaagtt |
| anti-MicF 33-16 | atgacggtaataaataaa |
| anti-MicF 33-18 | atgacggtaataaata |
| anti-MicF 33-20 | atgacggtaataaa |
| anti-MicF 33-22 | atgacggtaata |
| anti-MicF 33-24 | atgacggtaa |
| anti-MicF 16-3 | agttaatgatgata |
| anti-MicF 16-5 | agttaatgatga |
| anti-MicF 16-7 | agttaatgat |

Table S3: CPP-PNA sequences

| Name | Sequence[a] |
|---|---|
| Cell penetrating peptide (CPP) | KFFKFFKFFK |
| CPP-PNA scramble | KFFKFFKFFK-O-ttccgccaactc |
| CPP-PNA 12-1 | KFFKFFKFFK-O-aatgatgatagc |
| CPP-PNA 33-22 | KFFKFFKFFK-O-atgacggtaata |

[a]: the peptide or peptide-PNA conjugation sequences were shown from left to right following the N-terminus to C-terminus of polypeptide backbone

Table S4: Plasmids used in this study

| Plasmid # | Plasmid architecture | Name | Figure |
|---|---|---|---|
| MKT046 | J23119 – T1 (ColE1/KanR) | J23119 ColE1 control | 1B, 2, 4A, S1, S2, S3 |
| MKT050 | J23119 – MicF – T1 (ColE1/KanR) | J23119 MicF | 1B, 2, 4A, S1, S2, S3 |
| MKT053 | J23119 – ompF-sfgfp – T1 (p15A/AmpR) | J23119 ompF-sfgfp reporter | 1B, 2, 4A, S1, S2, S3 |
| MKT109 | J23119 – anti-MicF 33-1 – T1 (ColE1/KanR) | anti-MicF-33-1 | 1B |
| MKT131 | J23119 – anti-MicF 27-1 – T1 (ColE1/KanR) | anti-MicF-27-1 | 2 |
| MKT256 | J23119 – anti-MicF 24-1 – T1 (ColE1/KanR) | anti-MicF-24-1 | 2 |
| MKT138 | J23119 – anti-MicF 21-1 – T1 (ColE1/KanR) | anti-MicF-21-1 | 2 |
| MKT273 | J23119 – anti-MicF 18-1 – T1 (ColE1/KanR) | anti-MicF-18-1 | 2 |
| MKT244 | J23119 – anti-MicF 16-1 – T1 (ColE1/KanR) | anti-MicF-16-1 | 2, S1 |
| MKT165 | J23119 – anti-MicF 14-1 – T1 (ColE1/KanR) | anti-MicF-14-1 | 2 |
| MKT133 | J23119 – anti-MicF 13-1 – T1 (ColE1/KanR) | anti-MicF-13-1 | S1 |
| MKT294 | J23119 – anti-MicF 12-1 – T1 (ColE1/KanR) | anti-MicF-12-1 | 2, 4A |
| MKT329 | J23119 – anti-MicF 11-1 – T1 (ColE1/KanR) | anti-MicF-11-1 | S1 |
| MKT295 | J23119 – anti-MicF 10-1 – T1 (ColE1/KanR) | anti-MicF-10-1 | 2 |
| MKT121 | J23119 – anti-MicF 33-7 – T1 (ColE1/KanR) | anti-MicF-33-7 | 2 |

| | | | |
|---|---|---|---|
| MKT124 | J23119 – anti-MicF 33-10 – T1 (ColE1/KanR) | anti-MicF-33-10 | 2 |
| MKT258 | J23119 – anti-MicF 33-13 – T1 (ColE1/KanR) | anti-MicF-33-13 | 2 |
| MKT259 | J23119 – anti-MicF 33-16 – T1 (ColE1/KanR) | anti-MicF-33-16 | 2 |
| MKT277 | J23119 – anti-MicF 33-18 – T1 (ColE1/KanR) | anti-MicF-33-18 | 2 |
| MKT296 | J23119 – anti-MicF 33-20 – T1 (ColE1/KanR) | anti-MicF-33-20 | 2 |
| MKT142 | J23119 – anti-MicF 33-22 – T1 (ColE1/KanR) | anti-MicF-33-22 | 2, 4A |
| MKT267 | J23119 – anti-MicF 33-24 – T1 (ColE1/KanR) | anti-MicF-33-24 | 2 |
| MKT211 | J23119 – anti-MicF 16-3 – T1 (ColE1/KanR) | anti-MicF-16-3 | S1 |
| MKT192 | J23119 – anti-MicF 16-5 – T1 (ColE1/KanR) | anti-MicF-16-5 | S1 |
| MKT193 | J23119 – anti-MicF 16-7 – T1 (ColE1/KanR) | anti-MicF-16-7 | S1 |
| MKT220 | pLux – T1 (ColE1/KanR) | pLux ColE1 control | 1C, Table S5 |
| MKT284 | pLux – anti-MicF 33-1 – T1 (ColE1/KanR) | pLux-anti-MicF-33-1 | 1C, Table S5 |
| MKT327 | pLux – anti-MicF 12-1 – T1 (ColE1/KanR) | pLux-anti-MicF-12-1 | Table S5 |
| MKT221 | J23118 – T1 (pSC101/CmR) | J23118 pSC101 control | 3A, 4B, S4, S5A |
| MKT172 | J23118 – ompF-sfgfp – T1 (pSC101/CmR) | J23118 ompF-sfgfp reporter | 3A, 4B, S4, S5A |
| MKT116 | pLux – t500 (p15A/AmpR) | pLux p15A control | 3A, 4B, S4, S5A |
| MKT312 | pLux – MicF – T1 (p15A/AmpR) | pLux p15A MicF | 3A, 4B, S4, S5A |

Table S5: pLux-anti-MicF MIC values of individual replicates in μg/ml

| Antibiotic[a] | Construct | Rep 1[b] | Rep 2[b] | Rep 3[b] | Rep 4[b] | Avg[c] | p-value[d] | Figure |
|---|---|---|---|---|---|---|---|---|
| CEF | control | 64, 32 | 8, 16 | 16, 16 | 32, 16 | 25 | 0.0371 | 1C |
| | anti-MicF 33-1 | 16, 8 | 16, 8 | 8, 8 | 8, 8 | 10 | | |
| | control | 16, 32 | 64, 32 | 32, 32 | 16, 16 | 30 | 0.01333 | N/A |
| | anti-MicF 12-1 | 8, 16 | 8, 8 | 8, 8 | 8, 8 | 9 | | |
| NOR | control | 0.125, 0.125 | 0.125, 0.125 | 0.0625, 0.0625 | 0.125, 0.125 | 0.109 | 0.0335 | 1C |
| | anti-MicF 33-1 | 0.03125, 0.0625 | 0.03125, 0.03125 | 0.03125, 0.0625 | 0.03125, 0.0625 | 0.043 | | |
| | control | 0.125, 0.0625 | 0.125, 0.125 | 0.0625, 0.0625 | 0.0625, 0.0625 | 0.086 | 0.0568 | N/A |
| | anti-MicF 12-1 | 0.03125, 0.03125 | 0.0625, 0.03125 | 0.0625, 0.0625 | 0.0625, 0.03125 | 0.047 | | |

[a]: CEF, cephalothin; NOR, norfloxacin

[b]: two technical replicates were performed per biological replicate; technical replicate data is separated by a comma

[c]: average of all replicates (biological and technical)

[d]: A Wilcoxon Signed-Rank Test was performed between the control and anti-MicF conditions

Table S6: CPP-PNA MIC values of individual replicates in μg/ml

| Antibiotic[a] | Condition | Rep 1 | Rep 2 | Rep 3 | Rep 4 | Avg[b] | SD[b] | Figure |
|---|---|---|---|---|---|---|---|---|
| CEF | Scramble (6μM) | 16 | 16 | 16 | 16 | 16 | 0 | 3B, S5B |
| | 12-1 (6μM) | 16 | 32 | 8 | 16 | 18 | 10.1 | 3B |
| CEF | Scramble (8μM) | 8 | 8 | 8 | 8 | 8 | 0 | S5B |
| | MG1655 wildtype | 16 | 16 | 32 | 16 | 20 | 8 | S5B |
| CEF | Scramble (6μM) | 16 | 16 | 16 | 16 | 16 | 0 | 4C |
| | 33-22/12-1 (3μM each) | 8 | 8 | 8 | 8 | 8 | 0 | 4C |
| NOR | Scramble (6μM) | 0.0625 | 0.0625 | 0.0625 | 0.0625 | 0.0625 | 0 | 4C |
| | 33-22/12-1 (3μM each) | 0.0625 | 0.03125 | 0.0625 | 0.03125 | 0.0469 | 0.018 | 4C |
| NAL | Scramble (6μM) | 4 | 4 | 4 | 4 | 4 | 0 | 4C |
| | 33-22/12-1 (3μM each) | 2 | 2 | 2 | 2 | 2 | 0 | 4C |

[a]: CEF, cephalothin; NOR, norfloxacin, NAL, nalidixic acid

[b]: Average and standard deviation were calculated across the four biological replicates

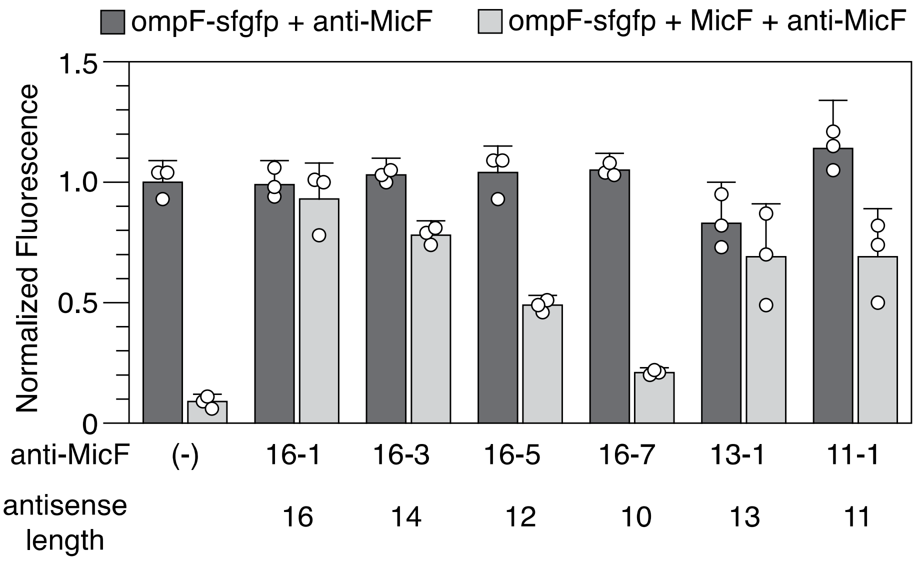


**Figure S1.** Screening anti-MicF asRNAs (continued). Average normalized SFGFP fluorescence of TX-TL reactions with and without anti-MicFs of various lengths. An anti-MicF is identified by the nucleotides of MicF that it was designed to target. Error bars represent standard deviation of three replicates.

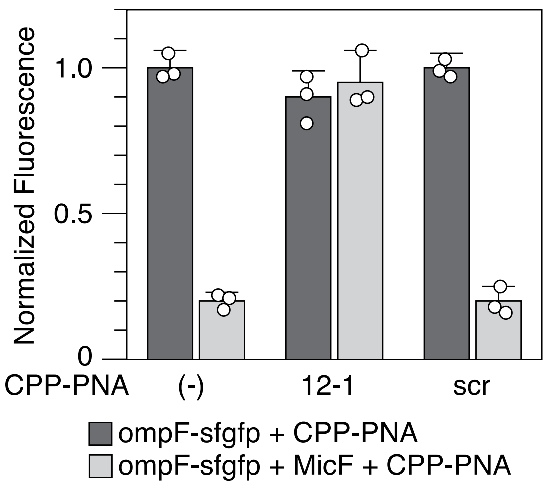


**Figure S2.** TX-TL test of CPP-PNA 12-1 and scramble. Average normalized SFGFP fluorescence of TX-TL reactions with and without CPP-PNA 12-1 or CPP-PNA scramble. Error bars represent standard deviation of three replicates.

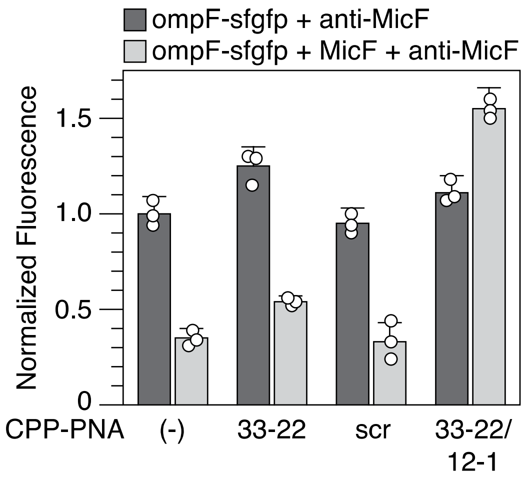


**Figure S3.** TX-TL test of CPP-PNA 33-22 alone and combination. Average normalized SFGFP fluorescence of TX-TL reactions with and without CPP-PNA 33-22 alone, CPP-PNA 33-22 in combination with CPP-PNA 12-1, or CPP-PNA scramble alone. Error bars represent standard deviation of three replicates.

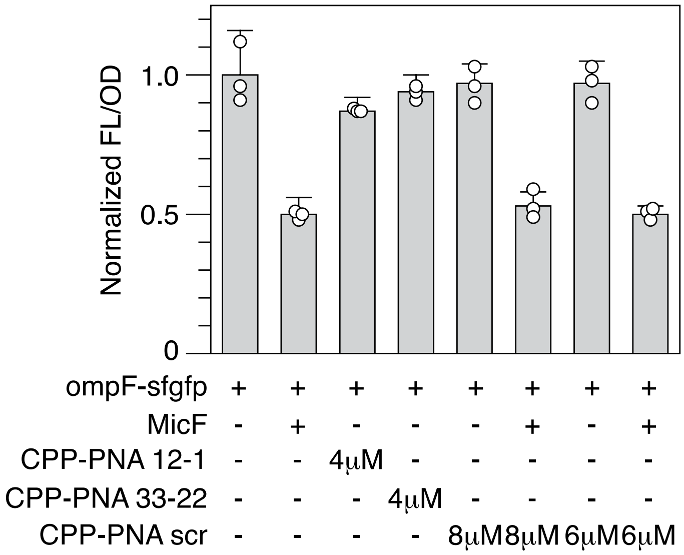


**Figure S4.** Controls for *E. coli* gene expression assay assessing CPP-PNAs. CPP-PNA 12-1, CPP-PNA 33-22, or CPP-PNA scramble was added to liquid cultures of *E. coli* MG1655 expressing the ompF-sfgfp reporter and an inducible MicF construct. AHL (2 nM) was added at the time of CPP-PNA addition to induce MicF expression. FL/OD was normalized to cells containing the ompF-sfgfp reporter alone. Error bars represent standard deviation of three biological replicates.

**A**

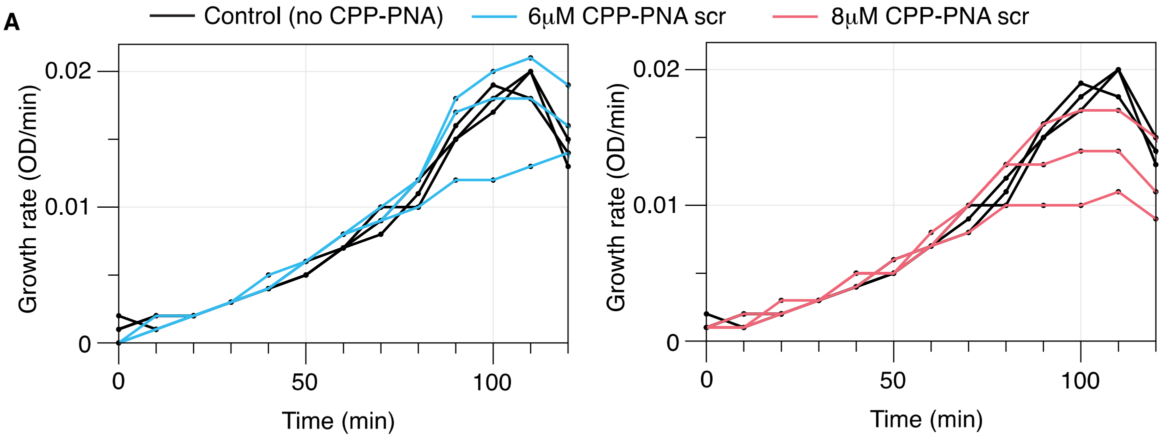


**B**

| | Average MIC (μg/ml) |
|---|---|
| | CEF |
| 6μM CPP-PNA scr | 16 |
| 8μM CPP-PNA scr | 8 |
| MG1655 | 20 |

**Figure S5.** Effect of CPP-PNA concentration on growth of *E. coli*. (A) Growth rate over time for cultures from Figure S4 with 6 or 8 μM of CPP-PNA scramble added to the culture. Each line represents a single biological replicate. Growth rates were compared to *E. coli* MG1655 with no added CPP-PNA. (B) MIC for cephalothin (CEF) for *E. coli* MG1655 with 0, 6, or 8 μM CPP-PNA scramble. Data represents the average of four biological replicates. Data for each replicate is found in Table S6.
